# *Amylase* copy number analysis in several mammalian lineages reveals convergent adaptive bursts shaped by diet

**DOI:** 10.1101/339457

**Authors:** Petar Pajic, Pavlos Pavlidis, Kirsten Dean, Lubov Neznanova, Erin Daugherity, Rose-Anne Romano, Danielle Garneau, Anja Globig, Stefan Ruhl, Omer Gokcumen

## Abstract

The amylase gene (*AMY*), which codes for a starch-digesting enzyme in animals, underwent several gene copy number gains in humans^1^, dogs^2^, and mice^3^, presumably along with increased starch consumption during the evolution of these species. Here we present evidence for additional *AMY* copy number expansions in several mammalian species, most of which also consume starch-rich diets. We also show that these independent *AMY* copy number gains are often accompanied by a gain in enzymatic activity of amylase in saliva. We used multi-species coalescent modeling to provide further evidence that these recurrent *AMY* gene copy number expansions were adaptive. Our findings underscore the overall importance of gene copy number amplification as a flexible and fast adaptive mechanism in evolution that can independently occur in different branches of the phylogeny.

## Introduction

Diet has been a significant evolutionary force in shaping human and nonhuman primate variation^4–6^. One of the best described examples of human-specific adaptation is the expansion of the copy number of the amylase gene in concordance with the increase of starch consumption in the human lineage^1^. A gene duplication in the ancestor of Old World monkeys and great apes initially led to the formation of two amylase genes (*AMY2A* and *AMY2B*) with pancreas-specific expression^7^. Then a subsequent gene duplication in the ancestor of great apes led to the formation of *AMY1* which gained salivary gland specific expression^8^. In the human lineage, further gene copy number gains of *AMY1*, but not *AMY2*, led to increased expression of the AMY1 enzyme in human saliva^1^. Copy numbers of amylase vary in different human populations^9^ and correlate with the extent of traditional starch consumption in these communities dating back only 10,000 - 20,000 years^1^. Despite all these gene copy number gains, which are thought to be mediated by non-allelic homologous recombination^1^, the coding sequences of the individual gene copies remained highly conserved. This suggests that maintenance of function was adaptively relevant.

While the evolution of the amylase locus in the human lineage is well described, the evolution of this locus in other mammals is less well understood. For example, it has been shown that mice, rats, and pigs express substantial levels of salivary amylase^10^. However, the evolutionary dynamics that led to gain-of-expression of amylase in saliva in these lineages remain unclear. Another interesting question is the evolution of amylase in domesticated animals. Recent studies have shown that dogs have also gained multiple copies of amylase after their split from wolves within only the last 5,000 years, likely as a result of their domestication^2,11^. As such, the evolution of amylase in other domesticated or human commensal mammals remains an alluring area of inquiry. Similarly, our understanding of the evolution of the amylase locus within the primate lineage remains limited. For example, it is not known why some Old World monkeys express substantial amylase activity levels in saliva, despite missing the great ape specific salivary amylase duplication^12^.

Here we address three areas of inquiry with regards to the evolution of the amylase locus in mammals: (i) Can the link between diet and amylase evolution, well-established in the human lineage, be generalized to other mammals? (ii) What are the evolutionary forces that shape amylase copy numbers in mammals? (iii) What are the genetic mechanisms leading to salivary expression in different nonhuman mammals? To answer these questions, we pursued a comprehensive investigation of amylase gene copy number and salivary expression across multiple mammalian lineages.

## Results and Discussion

### Recurrent amylase copy number gains in multiple mammalian lineages

The human-specific duplications of amylase are unique in their scope. Human genomes comprise up to 5 more haploid copies than chimpanzees. Moreover, most of these additional copies appear to contribute to expression of the amylase gene in saliva^1^. Therefore the recent revelation that a similar, independent, increase in amylase copy number occurred in dogs^2^ is remarkable, since it shows that the same gene independently underwent bursts of gene copy number gains in two separate species. To investigate whether these amylase copy number gains occur in other mammalian lineages as well, we conducted a digital droplet polymerase chain reaction (ddPCR) based analysis on amylase gene copy numbers from 153 DNA samples across 44 species encompassing all major branches of the mammalian phylogenetic tree. In addition to humans and dogs, we discovered similar bursts (*i.e.*, gain of more than one copy) of amylase gene copy number in house mice, brown rats, pigs, and boars (Figure 1, **Table S1**).

**Figure 1:**
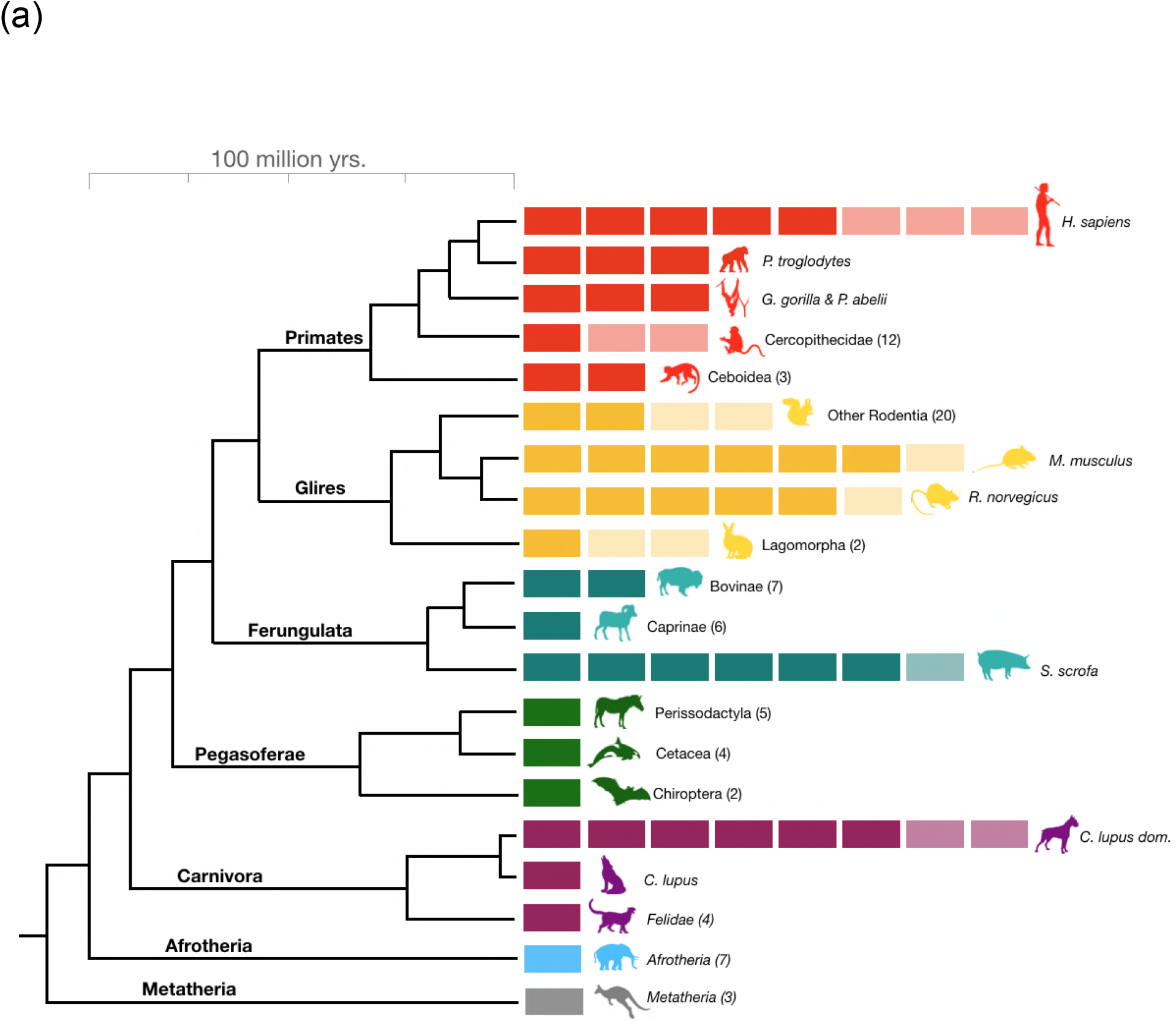
*Amylase* gene copy number bursts across mammals. Boxes represent haploid amylase gene copies among clades or of representative species across the mammalian phylogeny (see **Table S1** for a comprehensive dataset). Light-colored boxes represent the variation in copy numbers found in at least two individuals of a given species or in reference genomes of at least two species within a clade.

Given that copy number duplications occurred in different mammalian clades (Figure 1), we hypothesized that these events are a result of convergent evolution. Another possible explanation would be that the ancestor of placental mammals had multiple copies of the amylase gene, which were subsequently lost in particular mammalian lineages. To distinguish between these two scenarios, we constructed a maximum likelihood tree of amylase coding sequences from available reference genomes (Figure 2A). Our results showed that amylase genes within a given species are more similar to each other than they are to those of other species, suggesting that the duplication of amylase genes occurred independently in each lineage.

**Figure 2:**
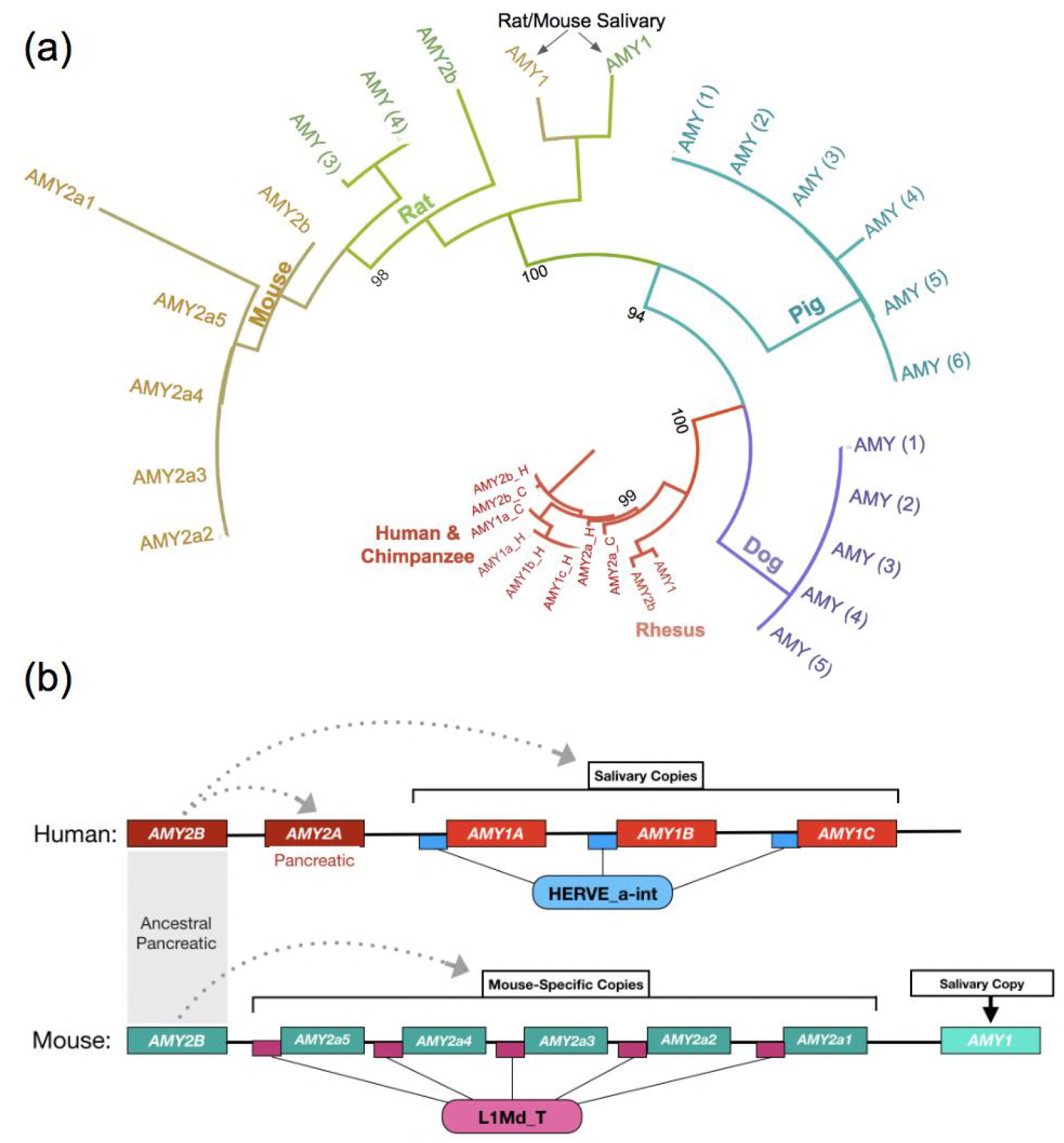
*Amylase* duplications evolved recurrently. **(a)** Maximum likelihood tree constructed using amylase protein sequences translated from reference genomes. **(b)** Depiction of the retrotransposons linked with amylase copies in mouse and human genomes. Small boxes symbolize the positions of mobile elements, HERVE_a-int LTR for humans (blue) and L1Md_T for mouse (purple). The dotted arrows indicate the likely origin of derived gene duplicates.

Samuelson *et al*. previously reported that a retrotransposon (HERV_a_int) was inserted upstream of a new amylase gene duplicate (*AMY1*) in the ancestor of great apes^7^. This copy rapidly duplicated several times in humans, carrying along the retrotransposon^1^. Based on this, we asked if a similar signature accounts for the copy number burst found in the mouse genome. We chose the mouse because its reference genome is adequately complete for such an analysis. Indeed, we found a mouse-lineage-specific retrotransposon (L1Md_T) in the upstream region of 5 out of the 7 mouse amylase genes. The presence of the retrotransposon along with the duplicated copies parallels the situation in humans (Figure 2B). Since different retrotransposons accompanied the rapid gene copy number gains in humans and mice, we conclude that these bursts occurred independently and, thus, are potentially a result of convergent evolution.

By ddPCR analysis, we found 9-13 diploid copies of the amylase gene in brown rats (Table S1). Considering the close phylogenetic relationship of rats and mice, we expected that the high copy number of amylase had evolved in their rodent ancestor. However, the L1Md_T retrotransposon is mouse-lineage specific. Therefore, the duplications in rats likely occurred independently from those in mice. We also confirmed the previous observations that dogs have gained at least 5 haploid copies of this gene over the short span of 5,000 years since their divergence from the wolf^11^. A similar process can be predicted for the pig and boar, whose genomes harbor 9-15 diploid copies of the amylase gene based on our analysis. In sum, our results suggest that amylase gene copy number gains have occurred recurrently in multiple, sometimes closely related, mammalian lineages.

### Amylase expression in saliva was facilitated through recurrent gene copy number gains independently in different mammalian lineages

Ancestral form of amylase in mammals codes for a pancreatic enzyme. However, in certain mammalian species, amylase also became expressed in saliva^13^. In humans, this acquisition of salivary gland-specific expression has been well documented^14^. It has been shown that the aforementioned retrotransposon insertion along with the *AMY1* duplicate in the ancestor of great apes is responsible for tissue-specific expression of this gene in salivary glands^7^. Previous studies also hypothesized that an independent, but similar gene duplication event led to the salivary expression of amylase in mice^8^. It remains unresolved whether the mechanism that enabled expression of amylase in mouse saliva is similar to that determined for humans. Moreover, even though various reports showed salivary expression of amylase in different mammalian species^12^, a comprehensive and systematic analysis of salivary expression of amylase across the mammalian clade is still missing.

To fill these gaps in knowledge, we performed a screen across the mammalian phylogeny to investigate which lineages express amylase activity in saliva. We used a two-pronged approach, comprising a starch lysis plate assay (Figure 3A) and a high-sensitivity in-solution fluorescence-based assay (Figure 3B). This approach provides the most comprehensive documentation of salivary amylase activity in mammals, encompassing 118 saliva samples across 20 species (Table S1). This is a significant contribution given that previous studies varied considerably in sample preparation, methods of analysis, and sensitivity^12^.

**Figure 3:**
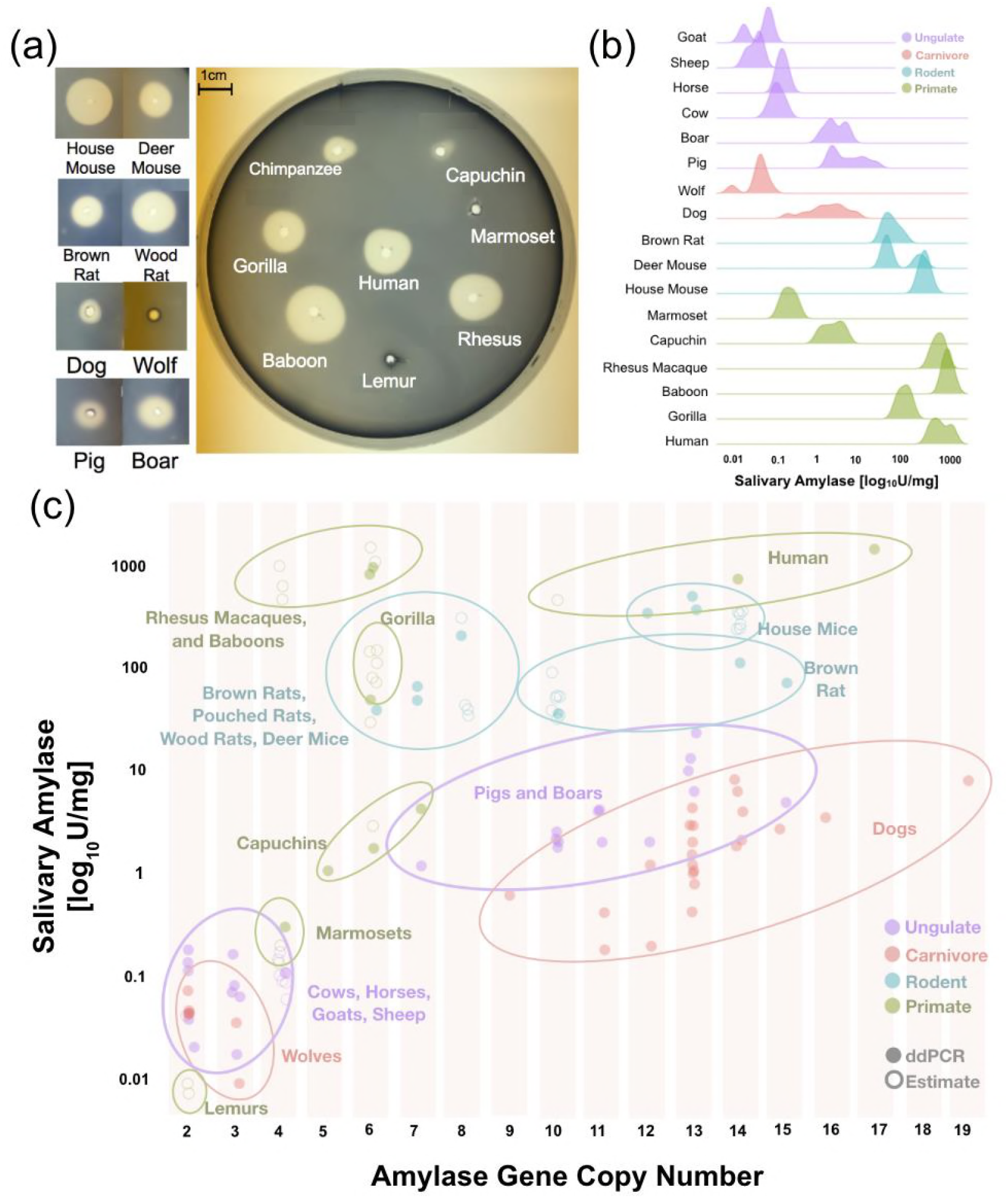
Salivary amylase activity and relationship to gene copy number. **(a)** A representative starch lysis assay plate showing the activity levels of amylase in the saliva of various mammalian species. **(b)** Density plots showing salivary amylase activity in different species. Full dataset can be found in Table S1. **(c)** Correlation of amylase activity and gene copy number in multiple different species. Data obtained by direct genotyping are represented by filled circles, while data estimated from reference genomes or through genotyping of other samples from the same species are represented by empty circles.

Our results showed that amylase activity in saliva is more widespread among mammals than previously thought (Figure 3B). In addition to species that were already known to express amylase in their saliva, we observed salivary activity in boars, dogs, deer mice, woodrats, and giant African pouched rats (Table S1). It is important to note here that our findings also suggest that salivary amylase activity in dogs varies from breed to breed (**Figure S1, Table S1**).

We surmised two competing scenarios to explain the observation that multiple mammalian lineages express amylase in their saliva. First, there could be independent gains of amylase expression in saliva spanning multiple lineages. Second, salivary expression of amylase could be an ancestral trait that was subsequently lost in most species. The above-described independent evolution of amylase gene copies in humans and mice supports the former hypothesis.

To further investigate this, we asked which of the mouse amylase copies is expressed in salivary glands by mapping available parotid salivary gland RNA-Seq data^15^ to the mouse reference genome (mm9) (Figure S2). We found that the copy annotated as mouse *AMY1* (Figure 2) is expressed in salivary glands, and is likely responsible for salivary expression of amylase in mice, while the other amylase duplicates have a negligible expression in salivary gland tissue (Figure S2). Mouse AMY1 has an amino acid sequence distinct from the other amylase copies in the mouse genome. This distinct sequence is shared with rats and other rodents (e.g., deer mouse, vole, mongolian gerbil, golden hamster), indicating that the duplication event that led to formation of *AMY1* likely has occurred in an ancestor of muroidea.

Even though more work will be needed to understand the regulatory mechanisms through which amylase gained salivary expression in pigs, boars, dogs, multiple rodents, and some Old World monkeys, it seems gene duplication is the required initiating step. Indeed, we found that the overall amylase gene copy numbers in species correlate well with observable enzymatic activity in saliva (Figure 3C). In fact, we could not find a species that underwent a “burst” of amylase gene copy number that did not show concurrent salivary amylase activity. Importantly, previous studies surmised that dogs do not express salivary amylase^2^, while we show here that several dog breeds express substantial amounts of this enzyme (Figure S1). This variable expression of amylase in saliva among different dog breeds makes this species an ideal model to study the mechanism of gain-of-expression in a new tissue facilitated by gene duplication. Overall, we conclude that the salivary activity of amylase has recurrently evolved in multiple mammalian lineages through gene duplication, where one or more of the duplicates have gained salivary gland expression.

### Varied diets correlate with increased amylase copy number

For humans, it has been postulated that starch consumption exerted a positive adaptive force on maintaining high amylase copy numbers^1^. Furthermore, the rapid copy number increase in dogs has been associated with their change in diet during domestication^2^. Based on these previous studies, we hypothesized that gains in copy number and the associated gain of amylase expression in saliva are likely driven by starch consumption. When we compared the amylase copy numbers in mammals that consume specialized diets (strict carnivores and non fruit eating herbivores) to those with broad-ranged diets, we found that the latter harbor significantly higher copy numbers of the amylase gene (**p=2.1×10^−7^, Mann-Whitney Test**, Figure 4A). We also found that the species consuming broad-ranged diets express significantly higher salivary amylase activity than those consuming specialized diets (**p=5.5×10^−4^, Mann-Whitney Test**, Figure 4B).

**Figure 4:**
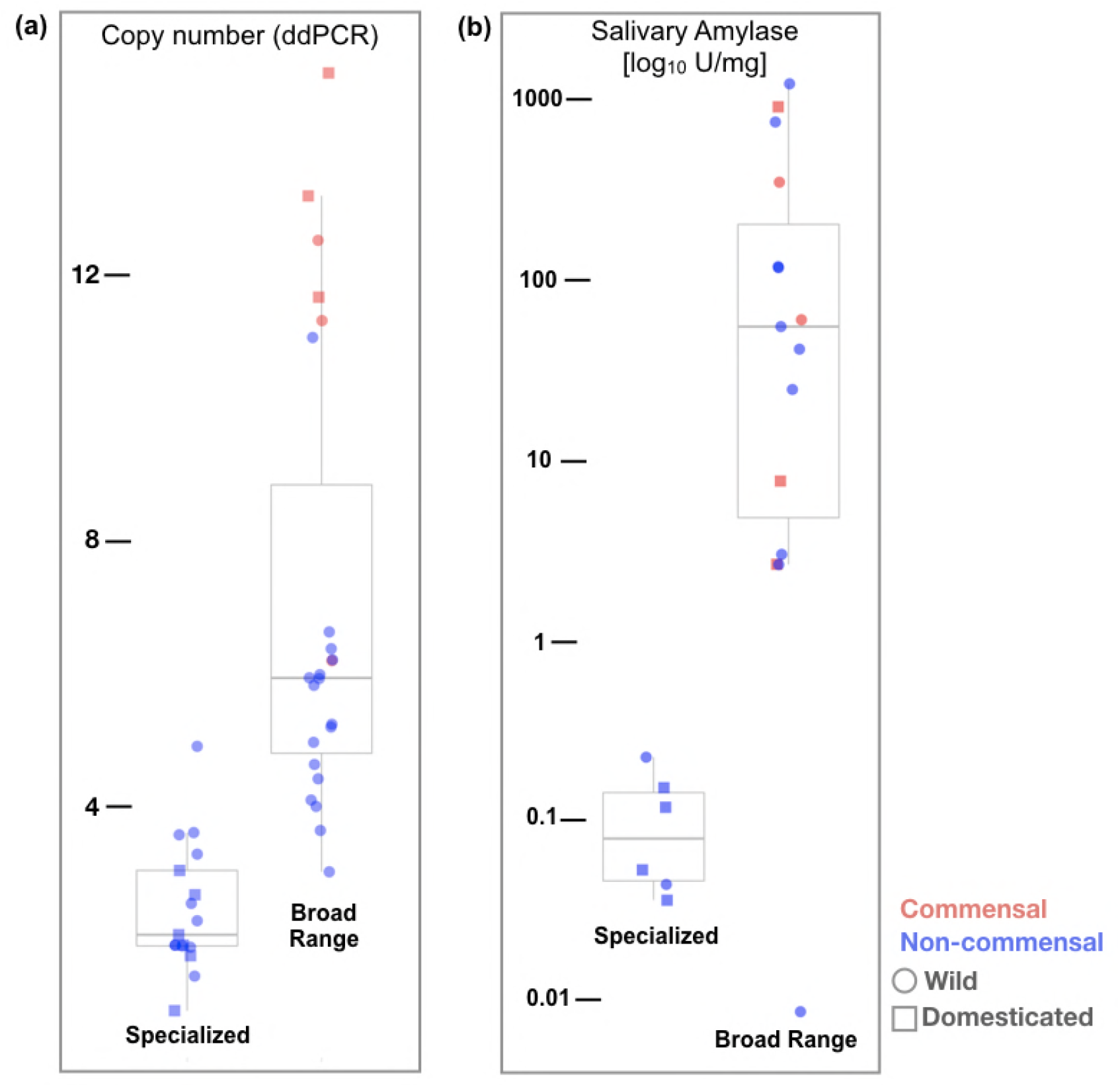
*Amylase* gene copy numbers and salivary enzyme activity correlate with diet. Box plot representing **(a)** *AMY* gene copy numbers or **(b)** salivary amylase activities in mammalian species assigned by their major diet. These include either as a specialized (carnivore or herbivore) or broad ranged diet. Dots and squares represent wild and domesticated species, respectively. Species that thrive in a commensal relationship with humans are shown in red while all others are shown in blue.

We then asked whether starch consumption is the main driver of the copy number gains and salivary expression of amylase. Unfortunately, there is no systematic survey of starch consumption among mammals, and the diet varies among subspecies, and even among populations of the same species^16^. As such, we could not reliably assess whether starch consumption by itself explains the copy number variation and salivary expression of the amylase gene. However, among all the species that consume a broad-ranged diet, we found that those who over recent evolutionary time have gained access to abundant starch-rich foods - either through domestication (as in the case of dogs and pigs) or through dietary commensalism with humans (as in the case of house mice as well as brown and black rats) harbor significantly higher copy number of the amylase gene (**p=1.2 × 10^−4^, Mann-Whitney Test**, Figure 4A). For salivary expression of amylase, this difference was not significant. This could potentially be due to the fact that most, if not all the species that consume a broad-ranged diet also consume starch to varying degrees.

Next, we conducted a comparative investigation of amylase copy number and its salivary expression between human-interacting species and their closest evolutionary relatives in the wild. In dogs, which due to their commensalism with humans consume a higher amount of starch than wolves, we noted a substantial increase over its ancestral state, not only in amylase gene copy number^2^, but also in salivary expression of amylase (Figure 3C, Figure S1). This increase was found less substantial in species that already consumed starch in their ancestral state (e.g. mice and rats which are granivorous). Along the same lines, we found no difference between domesticated pigs and wild boars. This could be explained because boars already consumed starch in amounts comparable to those of pigs. In fact, previous observations showed that boars and humans have similar starch-rich ancestral diets due to their consumption of underground starch-containing storage stem tissues known as tubers^17^.

### Evolution of amylase in primates

To understand how the broader trend of amylase evolution is reflected in the primate phylogeny, we have investigated multiple primate species, both for amylase gene copy number and salivary amylase activity (Figure 5). We confirmed previous studies which documented a duplication of the amylase gene in the ancestral population of the catarrhini and another duplication in the ancestral population of the great apes^8^. Among Old World monkeys, we found additional amylase gene copies in rhesus macaques, baboons, and vervets. In contrast, we found no additional gene duplication in leaf-eating old world monkeys (colobus, snub-nose and proboscis monkeys)^18^. Most New World monkey genomes that we tested carry 4 diploid amylase copies. Assuming that the ancestral state of this lineage had 2 copies, our results suggest another instance of gene copy number gain in the ancestor of New World monkeys. Moreover, we found an additional amylase copy in the capuchins, which consume more starch than other New World monkeys^19,20^. Next, we investigated lemurs, an outgroup primate species to monkeys and great apes, and found that they indeed only harbor 2 diploid copies of the amylase gene (Figure 5). This result in the lemur lineage, combined with the previous reports that ancestors of simians have a single copy^7,21^, suggest that primate ancestors had only one haploid copy of the amylase gene.

**Figure 5:**
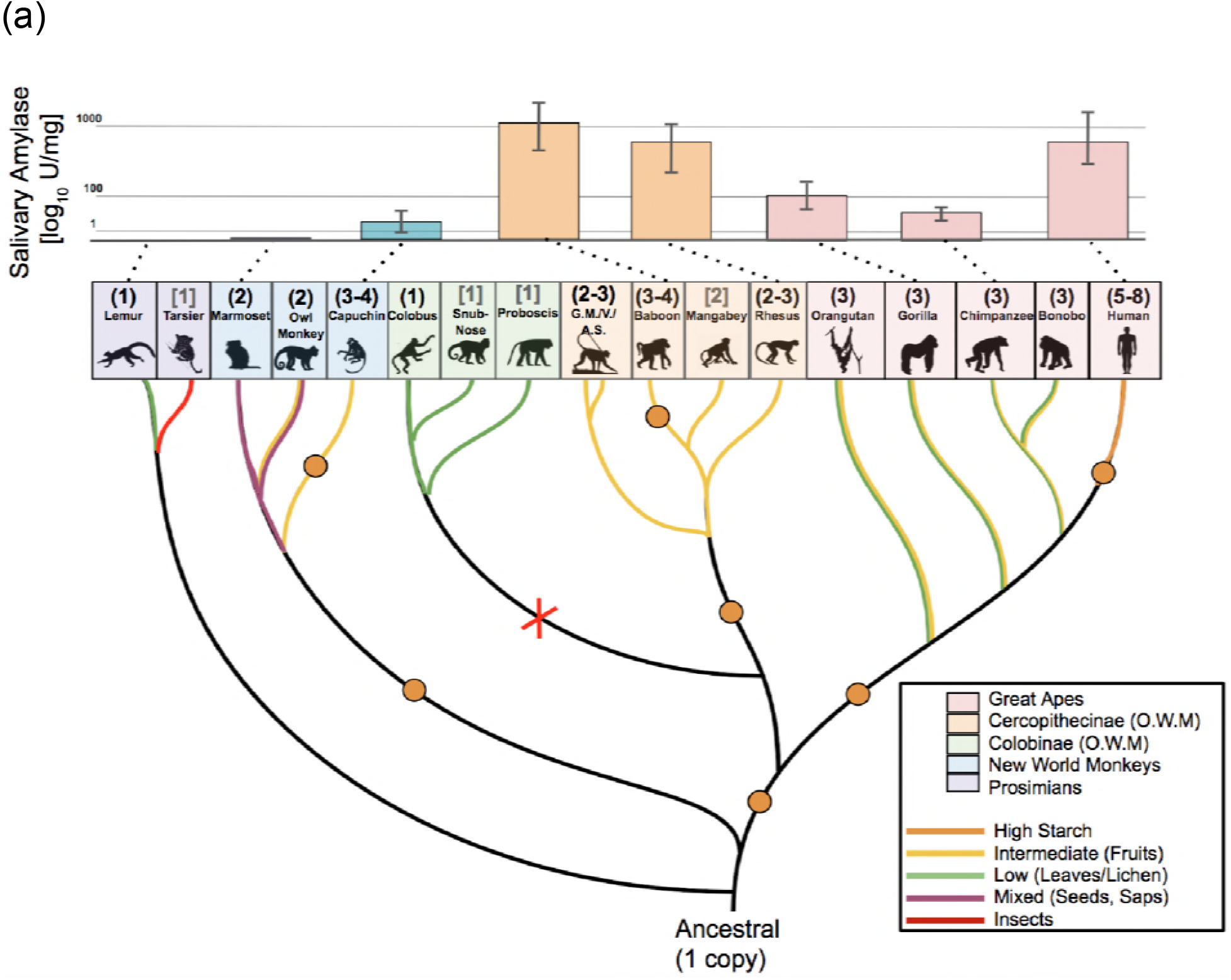
Amylase duplication events and salivary activities in the primate phylogeny. Bars represent mean amylase activity levels. Orange dots in the branches of the phylogenetic tree show independent duplication events of the amylase gene. The phylogeny represents the panel of primates we have obtained data for by ddPCR (*AMY* copy number in parentheses) or reference genome database information (copy number, grey, in brackets). The red X indicates an assumed copy number loss. Phylogenetic branches are colored according to diet preferences (see boxed insert). Abbreviations: G.M., green monkey; V, vervet; A.S., Allen’s swamp monkey.

Next we investigated whether variation in amylase gene copy numbers among primates translates into salivary expression, as we have shown for nonprimate mammals. We found that several species of Old World monkeys, including rhesus macaques and baboons, express abundant salivary amylase (Figure 5). These primates are known for their cheek pouches to store food for prolonged oral predigestion^22^, and previous studies have documented salivary activity of amylase in baboons^23^. New World monkeys consume even more diverse diets than Old World monkeys. For example, marmosets primarily consume insects and plant exudate^24^, while owl monkeys consume flowers, insects, nectar, and leaves^20,25^. Capuchin monkeys consume fruits, bulbs and seeds^19,20^. In agreement with these dietary habits we found little or no salivary activity of amylase in New World monkeys. The only exception were capuchins, which we discovered to express salivary amylase, and which also consume a higher proportion of starch in their diet compared to the others (Figure 5).

Combined, our results in primates document additional instances where lineage-specific duplications of the amylase gene in the cheek pouched cercopithecines and capuchins coincide with salivary expression. Broadly, our results suggest that the evolution of the amylase locus in primates follows the general trends observed for all mammals in that dietary strategies rapidly shape both the copy number and salivary expression in a lineage specific manner.

### Modeling the evolution of amylase copy number

Our empirical analyses of amylase copy number variation across mammals clearly show a trend where animals consuming high amounts of starch, carry higher copy numbers of this gene (Figure 4A). This aligns well with the hypothesis that high amylase copy number is adaptively maintained in these lineages^8^. To formally test this hypothesis, we simulated the copy number in 100 animal species (available through Hg19 100way conservation alignment^26^) under the assumption of neutrality (see Methods section) (**Table S4, Figure S3**). In our simulations none of the neutral models could explain the observed copy number variation in the amylase locus. On one hand, simulations under higher mutation rates could not explain the observation that certain distantly related mammalian lineages such as humans, dogs, pigs, mice, and rats harbor similar amylase copy numbers. While on the other hand, simulations under low mutation rates could not explain the observation that certain closely related species, such as humans and chimpanzees or wolves and dogs, harboring substantially different amylase copy numbers. Thus, this simulation-based analysis shows that the observed copy number variation among mammals cannot be explained by neutral evolution alone. In the light of the empirical analyses described in this study, we argue that the most parsimonious explanation is that lineage-specific, convergent adaptive forces have shaped copy number variation of the amylase gene among mammalian species.

## Conclusion

Our results reveal a staggering diversity of amylase gene copy numbers across extant mammals that correlates with starch consumption. We report multiple bursts of amylase copy number gains that occurred independently in different lineages. Furthermore, our results showed that each of these bursts led to expression of amylase in saliva, providing a case example of convergent evolution of gene regulation by structural variation in a diet-related gene.

Our results also raise intriguing questions that could not be resolved in this study: 1. How do putative salivary gland-specific enhancers evolve along with the gene copy number to lead to amylase expression in salivary gland tissue? 2. Is there any functional variation among amylase gene copies, either through sequence variation or differences in post-translational modifications? 3. Why and how can diet have such a dramatic adaptive effect on copy number of a gene, and what are the selective advantages gained by increased expression of amylase in saliva? Our results showed that phylogenetically distant species with diverse food preferences and habitats have evolved similar amylase gene copy numbers, which correlate well with known levels of starch consumption. This fits into an evolutionary explanation where increase in copy number leads to higher amylase expression, which in-turn allows rapid and effective intestinal digestion of starch.

We further showed that amylase is expressed in the saliva of species consuming a broad-ranged diet. Most mammalian species, including humans, primarily digest starch in their digestive tract rather than in the oral cavity. As such, a simple explanation based on digestion alone fails to fully explain the gain of salivary expression of this gene even in high starch-consuming species. Based on our results, we argue that such putatively adaptive expression of amylase in saliva depends on the ecological and behavioral context of the species and, thus, is lineage-specific. For example, it is remarkable to see the dramatic increase of salivary amylase activity in the cheek-pouched Old World monkeys, which conduct almost half of their starch digestion in their oral cavity. In other species, food is not retained long enough in the mouth for substantial starch digestion to take effect. Consequently, indirect effects of salivary amylase activity other than solely digestion may also play a role in how natural selection acted on the regulation of this gene. In this context, other studies found links between salivary amylase and taste perception^27^, metabolic regulation^28^, and bacterial composition in the oral cavity^29,30^. Overall, one can argue that presence of amylase enzymatic activity in saliva may shape food preference and even niche partitioning among omnivorous mammals living in starch-rich ecologies, followed by coevolution with the oral microbiome.

## Methods

### Samples

We chose our panel of mammalian species based on their phylogeny, diet preference (carnivore, herbivore, omnivore), domestication, and commensal relationship with humans.

Overall we compiled 153 DNA samples from 44 different species and 118 saliva samples from 20 different species. Detailed information about the samples used in this study and their sources can be found in Table S2. The diet information for individual species was mostly acquired from Michigan Animal Diversity Web (https://animaldiversity.org/), unless other more specific studies were cited.

### Genomic analysis

DNA was isolated from buccal swabs and saliva using a commercially available kit (ChargeSwitch^®^ gDNA Buccal Cell Kit, Invitrogen). DNA extraction from blood and cell lines was conducted as described previously^31^. The DNA was analyzed by digital droplet PCR (ddPCR) to determine amylase gene copy number. For primer design we targeted amylase exonic sequences that are conserved among copies and between species. The primer sets used for each species are listed in **Table S3**. In most species, ddPCR results were highly concordant with copy number estimations based on BLASTx and BLASTp analysis (**Figure S4**). Only in certain species, disparities between our ddPCR results and existing databases were noted (**Table S1**, Figure 3C).

### Phylogenetic analysis

Amino acid sequences translated from reference genomes for the amylase gene copies were downloaded from NCBI. Sequences were aligned and a phylogenetic output was generated using a custom *Python* code as described previously^32^. We constructed a maximum likelihood tree from the protein sequences using RAxML^33^, bootstrapping with 1000 replicates for branch support. Visualization was performed using FigTree^34^.

### Measurement of amylase enzymatic activity

We used two methods to measure salivary amylase activity. First, we conducted a direct measurement of enzyme activity using a starch lysis agar plate (Figure 3A) following a previously described protocol^35^. In parallel, we used a high-sensitivity (detection limit 2 × 10^−3^ U/ml) microtiter plate assay (EnzCheck *Ultra* Amylase Assay Kit, Invitrogen) following the manufacturer’s protocol and using α-amylase from human pancreas (Sigma) as the standard. Total protein concentrations were measured using the bicinchoninic acid (BCA) assay (micro-BCA, BioRad) with bovine serum albumin as the standard. Optical density measurements were performed using the Nanodrop 2000 spectrophotometer (Thermo Fisher).

### Simulations

We simulated the neutral intra- and inter-species copy number variation in 100 animal species using the software CoMuS^36^ and the phylogenetic tree provided by the UCSC Genome Browser (http://hgdownload.cse.ucsc.edu/goldenpath/hg19/multiz100way/). The original version of CoMuS performs neutral multi-species coalescent simulations, thus it separates the coalescent process from the mutation process. This assumption, however, may be inappropriate for studying the evolution of copy number since the mutation rate at a specific lineage at time *t*, may depend on the present copy numbers on this lineage at time *t*. For example, on the one hand, a large number of copies present may imply an increased mutation rate. On the other hand, a small number of copies present may result in a decreased mutation rate. At the extreme, zero copies represent an absorbing state, i.e. no further changes are possible. Also, for a single copy, a reasonable assumption is that a gain should occur more frequently than a loss. Such assumptions related to the neutral copy number evolution result in a dependence of the mutation rate on the pre-existing copy number state. Thus, we implemented a modified version of CoMuS, where genealogies are simulated first, and thereafter mutations occur along the branches using a pre-order traversal of the tree: each mutation may affect the mutation rate on each subtree that has inherited it. We simulated neutral copy number variants for a total of 300 individuals, that is 3 individuals for each of the 100 species of the guide phylogenetic tree (**Table S4**). The modified version of CoMuS that we used here can be downloaded from https://github.com/idaios/comuscnv.

## Data analyses and figures

All the input data are provided in **Tables S1** and **S4**. We used custom scripts to analyze data and produce the figures primarily using the R statistical package.

## Acknowledgements

We would like to recognize and thank all the individuals and institutions who provided us with DNA and saliva samples from different mammalian species: Joseph Cook and Mariel Campbell (Museum of Southwestern Biology, Division of Genomics Resources, University of New Mexico), Coriell Institute for Medical Research, Karen Davis (Wolf Park, Battle Ground, IN), Barbara McCabe (SUNY at Buffalo), Luce E. Guanzini (Center for Animal Resources and Education, Cornell University), Klaus Depner (Friedrich-Loeffler-Institut, Greifswald, Germany), Ann-Marie Torregrossa (SUNY at Buffalo; Department of Psychology), Michele Skopec (Weber State University; Department of Zoology), Jill Kramer (SUNY at Buffalo; Department of Oral Biology), Kurt Volle, Alicia Dubrava and fellow gorilla caretakers of the Buffalo Zoo, the Southwest National Primate Research Center funded by NIH grant ORIP/OD P51OD011133, the Yerkes National Primate Research Center which is funded by NIH grant ORIP/OD P51OD011132), and the Duke Lemur Center funded by NSF grant LSCBR #1050035.

We would also like to thank Ozgur Taskent, Jessica Poulin, Trevor Krabbenhoft, and Derek Taylor for careful proofreading of this manuscript and discussions to better present the data and Aggelos Koropoulis for the implementation of CoMuSCNV, This study was funded by National Institute of Dental and Craniofacial Research grants DE019807 and DE025826 (SR).

## Supplementary Figures

**Figure S1: Dog copy number versus salivary amylase activity.** X-axis represents the haploid copy number of various dog breeds, (see Table S1 for breeds). Y-axis represents the salivary enzymatic activity for the same sample. A trendline was applied to show correlation. Red dots represent individual dog sample.

**Figure S2: RNA-sequencing for expression of amylase genes in mouse salivary gland.**

Green boxes on the x-axis represent the gene order on the mouse reference genome. The y-axis is drawn in log scale and represents the fragments per kilobase of exon per million reads (FPKMS) from RNA sequencing. The purple bars designate the average FPKMS read coverage for RNA from 2 adult mice (12 weeks of age) for their parotid salivary glands. The gene schematic diagram displays the RNA sequencing coverage across the exons of *AMY1*. Data were extracted from Gluck et al.^15^.

**Figure S3: Simulation results.** To visualize our simulated dataset, we plotted the phylogenetic distance between species in a pairwise fashion on the x-axis and the absolute copy number difference between species on the y-axis Phylogenetic distance is defined as the total length of the branches that separate two species. The phylogenetic tree was downloaded from the UCSC multiz100way file related to humans. The mutation rates for different simulations are noted near the lines. Mutation rates correspond to 4N_e_μ, where Ne is the effective population size, and N_e_ the mutation rate per locus and per generation. The upper graph shows simulations across all species contained in the multiz100way database (http://hgdownload.cse.ucsc.edu/goldenpath/hg19/multiz100way/). The bottom diagram focuses on the mammalian species under investigation in this study. The observed data (red dots) and the red fitted-line were superimposed on the simulation results.

**Figure S4: Reference Genome Accuracy.** The y-axis represents the haploid copy number data obtained from available references genomes. The x-axis represents our ddPCR copy number data for 27 different species. A linear regression line is plotted to visualize the correlation.

